# Decoding of Speech Information using EEG in Children with Dyslexia: Less Accurate Low-Frequency Representations of Speech, Not “Noisy” Representations

**DOI:** 10.1101/2022.05.02.490279

**Authors:** Mahmoud Keshavarzi, Kanad Mandke, Annabel Macfarlane, Lyla Parvez, Fiona Gabrielczyk, Angela Wilson, Sheila Flanagan, Usha Goswami

**Author notes:** MK: Conceptualisation, Methodology, Data analysis, Visualisation, Writing original draft. KM: EEG Methodology, Investigation, Writing – review and editing. AM: EEG Investigation, Behavioural Data Curation. LP: EEG Investigation. FG: Behavioural Investigation, Data Curation. AW: Behavioural Investigation, Data Curation. SF: Behavioural Investigation, Writing - review and editing. UG: Conceptualisation, Funding Acquisition, Methodology, Resources, Supervision, Writing Original Draft.

## Abstract

The amplitude envelope of speech carries crucial low-frequency acoustic information that assists linguistic decoding. The sensory-neural Temporal Sampling (TS) theory of developmental dyslexia proposes atypical encoding of speech envelope information <10 Hz, leading to atypical phonological representations. Here a backward linear TRF model and story listening were employed to estimate the speech information encoded in the electroencephalogram in the canonical delta, theta and alpha bands by 9-year-old children with and without dyslexia. TRF decoding accuracy provided an estimate of how faithfully the children’s brains encoded low-frequency envelope information. Between-group analyses showed that the children with dyslexia exhibited impaired reconstruction of speech information in the delta band. However, when the quality of speech encoding for each child was estimated using child-by-child decoding models, then the dyslexic children did not differ from controls. This suggests that children with dyslexia encode neither “noisy” nor “normal” representations of the speech signal, but different representations.

## 1. Introduction

Dyslexia is a disorder of development. Classically, a child has shown apparently typical language acquisition and cognitive development until faced with the task of learning to read. Suddenly, the affected child shows a specific problem with reading and spelling that cannot be accounted for by low intelligence, poor educational opportunities, or obvious sensory or neurological damage. Reading is the process of understanding speech when it is written down. To acquire early reading skills, the child must learn the visual code used by their culture for representing speech as a series of symbols. Logically, individual differences in acquiring reading could be related developmentally to either spoken language processing or visual code processing or both (Snowling, 2000; Stein and Walsh, 1997). However, it is important to recognise that the visual code that must be learned is not a neutral visual stimulus. It is a culturally-specific code that is taught directly using symbol-sound correspondences, such as the alphabetic correspondences used to represent English phonology (the sound system of the language). Learning this visual code typically begins a few years into the development of a spoken language system, and it is now widely recognised that individual differences in children’s pre-reading *phonological awareness* (their ability orally to recognise or manipulate phonological units in words such as syllables, rhymes or phonemes) is a causal determinant of how readily their visual code learning progresses (Ziegler and Goswami, 2005, for cross-language review). Indeed classically, ‘deficits’ in the representation and use of phonological information have been seen as critical in the etiology of developmental dyslexia (Catts, 1993; Stackhouse and Wells, 1997; Stanovich and Siegel, 1994; Swan and Goswami, 1997; Snowling, 2000).

To understand the nature of this ‘phonological deficit’ in dyslexia, longitudinal studies that begin with infants are required. Evidence from both electrophysiological and behavioural measures suggests that infants at family risk (genetic risk) for dyslexia already exhibit deficits in the auditory processing of speech and non-speech sounds in their first year of life, which could be expected to affect novel word learning. It is well known that the speech signal provides a rich repertoire of acoustic cues that infants exploit to learn new words (Saffran, 2001). Although very few neurophysiological longitudinal studies of at-risk infants exist, EEG studies suggest atypical detection of changes to the acoustic features of speech and non-speech sounds (e.g., F0 frequencies, vowel length, and consonant voice onset time) in newborns and two-month-olds, and there are also differences in their hemispherical distribution of the neural responses to these acoustic stimuli (Guttorm, Leppänen, Hämäläinen, Eklund, and Lyytinen, 2010; Guttorm et al., 2005; Leppänen et al., 2010; see also van Zuijen, Plakas, Maassen, Maurits, and van der Leij, 2013). Regarding the atypical development of phonological representations for words that characterise older children once they are diagnosed with dyslexia, the most commonly accepted hypothesis is that these phonological representations are somehow “noisy” or “imprecise” (Swan and Goswami, 1997; Snowling, 2000). As explained in detail by Ramus and Svenkovits (2008), the general idea concerning “noisy” representations is that a dyslexic child’s neural representations of the speech signal are somehow fuzzier, for example having a lower resolution than they should, or preserving too many acoustic or allophonic details (Adlard and Hazan, 1998; Elbro et al., 1998; Manis et al., 1997; Mody, Studdert-Kennedy, and Brady, 1997; Serniclaes et al., 2004; Bogliotti et al., 2008; Snowling, 2000; Serniclaes and Seck, 2018). Other terms used in the literature to describe dyslexic children’s phonological representations are “imprecise” or “under-specified”. Ramus and Svenkovits (2008) disputed this idea about “noisy” representations, arguing that the phonological representations of individuals with dyslexia may be quite normal. The phonological ‘deficit’ might arise instead from problems in *accessing* these normal representations when required for phonological awareness and other oral tasks. Although access is not mechanistically defined, Ramus and Svenkovits expect access to be more difficult when a task requires lots of phonological information to be stored in short-term memory, or a task requires speeded or repeated retrievals of phonological information, or it requires the extraction of speech information from noise (Ramus and Svenkovits, 2008). To our knowledge, however, neural studies have not yet been able to contrast the “noisy representations” view of the phonological ‘deficit’ in dyslexia with the opposing view of Ramus and Svenkovits that the phonological representations developed by the dyslexic brain are “normal”. We describe an initial neural approach to exploring this contrast here.

In the last 5 years, neural studies with children with dyslexia have been applying novel computational methods to EEG and MEG data to try to measure the quality of children’s phonological representations directly. These methods either use original speech features such as the envelope to predict the brain’s neuroelectric responses (forward models, from speech envelope to brain response, Di Liberto et al., 2018), or reconstruct speech-based representations of envelope information from the brain’s neuroelectric responses to speech input (backward or speech reconstruction models, from brain response to speech signal, Power et al., 2016; Destoky et al., 2020). These studies use a linear approach to stimulus reconstruction or to neural prediction, namely temporal response functions (TRFs, Di Liberto et al., 2015; Crosse et al., 2016). Backward TRF models reconstruct either the broadband speech envelope in the original signal or the envelopes in different frequency bands (such as 0.5 – 1.5 Hz or 2 – 8 Hz; Destoky et al., 2020). Such studies are still rare in the literature, and only one of them to date has used natural speech (Di Liberto et al., 2018) rather than degraded speech (vocoded speech, Power et al., 2016) or speech-in-noise (Destoky et al., 2020). Nevertheless, all studies to date show atypical low-frequency encoding of the speech signal by children with dyslexia, with speech envelope information <10 Hz encoded reliably less accurately by dyslexic children compared to age-matched control children.

These prior TRF studies also included younger children matched for reading level to the children with dyslexia. This is an important experimental control for the effects of learning on the developing brain. It is known that learning to read in itself changes phonological representations, as orthographic knowledge gained through reading affects performance in oral listening tasks for both children and adults (Ehri and Wilce, 1980; Ziegler and Ferrand, 1998). In principle, younger reading-level-matched (RL match) children provide a control for the effects of reading experience on the developing language system. Both of the prior EEG studies using both backward and forward TRF models reported significantly greater stimulus reconstruction accuracy (Power et al., 2016) or neural response accuracy (Di Liberto et al., 2018) regarding low-frequency acoustic speech information for their RL controls compared to their dyslexic participants. Accordingly, in EEG studies the dyslexic brain shows less accurate speech encoding of low-frequency envelope information than the brains of younger children matched for reading experience, suggesting a fundamental difference in encoding certain aspects of linguistic information. The exception was the MEG study using speech-in-noise tasks, and the authors explained their null result by suggesting that discriminating speech in noise may be facilitated by learning to read (Destoky et al., 2020). If speech-in-noise performance depends on the level of reading attained, then children with dyslexia should perform at the same level as younger children matched for reading, as found by Destoky et al. (2020).

To date, therefore, data from neural TRF studies do not suggest that the phonological representations of words developed by children with dyslexia are “normal” regarding the encoding of low-frequency acoustic information. At the same time, however, it is not clear that these representations are “noisy” or “fuzzy” regarding this low-frequency information. Rather, low-frequency acoustic information in the speech signal appears to be represented *less accurately* in the neural mental lexicons of children with dyslexia. This suggests that speech itself may be perceived differently by children with dyslexia. Rather than being a noisy signal for the dyslexic brain, the signal itself may be represented in an atypical manner, with less accurate representation of low-frequency information (Goswami, 2022). This reduced accuracy for envelope information may possibly be compensated by an over-weighting of allophonic and other information (Bogliotti et al., 2008), which may be represented with greater specificity by children with dyslexia. It is also possible that both low-frequency envelope information in the speech signal and other speech features like voicing are represented inaccurately by the brains of children with dyslexia.

These potential differences in the encoding of different types of acoustic information can be assessed in part with psychophysical methods (Serniclaes et al., 2004; Goswami et al., 2011; Serniclaes and Seck, 2018). For example, Goswami et al. (2011) presented a phonetic contrast (ba/wa) to children with dyslexia and age-matched and RL controls, changing “ba” to “wa” either by varying amplitude rise time in synthetic syllables or by varying frequency rise time. The children with dyslexia were significantly poorer at discriminating “ba” from “wa” when the phonetic change depended on amplitude rise time, but were significantly *better* than both age-matched and RL controls when the phonetic change depended on frequency rise time. These differences in the encoding of acoustic information appear to impair dyslexic children’s performance in classical phonological awareness tasks while enhancing their performance in some speech-based psychophysical tasks such as categorical perception (Serniclaes et al., 2004; Bogliotti et al., 2008). These representational differences regarding speech information would also complicate the process of learning to read, since any visual symbol system that is being learned will have been designed for learners who hear speech differently from children with dyslexia. This would apply whether the visual code is an alphabetic system or any other orthographic system (see Goswami, 2022).

In the current study, we set out to use a stimulus envelope reconstruction approach to begin to tackle these theoretical issues (“noisy” representations versus “normal” representations) with the same data. Our reasoning was as follows. Using a backward TRF modelling approach, thereby using neurophysiological responses to reconstruct speech envelopes < 10Hz, natural speech listening data can be used either to compare the decoding accuracy of the models that are reconstructed between groups of children, or within individual children. Envelope decoding accuracy can be taken as an estimate of how faithfully the children’s brains are encoding low-frequency envelope information present in the speech signal. If children with dyslexia are developing phonological representations that do not encode low-frequency acoustic information as accurately as control children, then the between-group comparison should reveal significant differences in the accuracy of the decoding models. In such a case, the phonological representations developed by the children with dyslexia could not be designated as “normal”. If children with dyslexia are encoding “noisy” or “fuzzy” representations of the speech signal, then the within-child comparison should also reveal significant differences in the accuracy of the models at the group level. The averaged within-child models of low-frequency envelope information for the children with dyslexia should be less accurate than the averaged within-child models for the control children, as the speech-based representations of this low-frequency information developed by children with dyslexia should be less consistent. To investigate these possibilities, EEG was recorded while children listened to a 10-minute story presented as audio-only. Backward TRF models were then computed, and the accuracy of these models was compared for the canonical delta, theta and alpha (control) bands using both between-group and within-child comparisons. The alpha band was utilised as a control band, as Temporal Sampling (TS) theory (Goswami, 2011) would not predict group differences for this band. In adults, alpha band oscillations are mostly related to working memory and attention, but some studies have suggested that the alpha band plays an important role in auditory processing as well regarding continuous speech perception (Strauß et al., 2014; Dimitrijevic et al. 2017). In the adult literature, cortical activity in the delta band is mostly associated with prosodic, intonational and phrasal features of speech (Ding and Simon, 2014), while cortical activity in the theta band helps to identify the onsets of syllables, contributing to speech parsing (Ding and Simon, 2014; Di Liberto et al., 2015; Keshavarzi et al., 2020; Keshavarzi and Reichenbach, 2020). For the between-group comparisons, we expected to replicate the finding in the literature that low-frequency acoustic speech information is encoded less accurately by children with dyslexia in the delta and theta bands. Regarding the “noisy” representations question, given the frequent observation that children with dyslexia appear to show no difficulties in speaking and listening tasks that do not tax phonological knowledge, it may be that the children are operating with perceptually stable phonological representations at the speech envelope level that are not “noisy” for their users.

## 2. Material and Methods

### 2.1. Participants

Fifty-one children were participated in this study. Twenty-one participants were typically developing children (mean age of 109.3 ± 5.4 months) and thirty participants had developmental dyslexia (mean age of 110.7 ± 5.6 months). The unequal group sizes arose due to Covid-19, which necessitated the cessation of testing part-way through the study, thereby also preventing testing of the recruited RL control group. One of the dyslexic children only completed 5 minutes of the EEG session, due to lots of head movement, not listening to the story, and taking out the earphone phone during data collection. We therefore excluded this child from further analysis. The children with dyslexia did not have any additional learning difficulties (e.g., ADHD, dyspraxia, autistic spectrum disorder, developmental language disorder), and were recruited through learning support teachers. The absence of the additional learning difficulties was confirmed based on school and parental reports and our own behavioural testing. All participants had a nonverbal IQ above 84, and their first language spoken at home was English. All children received a short hearing screen across frequency range of 0.25 – 8 kHz (0.25, 0.5, 1, 2, 4, 8 kHz) using an audiometer, and they all were found to be sensitive to sounds within the 20 dB HL range. SES data were not formally collected, but children were attending state schools (equivalent to US public schools) situated in a range of towns and villages near a university town in the United Kingdom. Participants and their parents gave informed consent for the EEG study in accordance with the Declaration of Helsinki, and the study was approved by the Psychology Research Ethics Committee of the University of Cambridge.

### 2.2. Behavioural tests

A series of standardised tests (see Table 1) of written and spoken language were administered in schools prior to the EEG session to assess cognitive development (please see Keshavarzi et al., 2022, for a full description of each test; the current participants also received the repetitive speech task employed in that study). Reading and spelling were assessed using the British Ability Scales (BAS, Elliot et al., 1996) and the Test of Word Reading Efficiency (TOWRE, Torgesen et al., 1999). Four subscales from the Wechsler Intelligence Scale for Children (WISC-V, Weschler, 2016) including two verbal (vocabulary and similarities) and two non-verbal (block design and matrix reasoning) scales were administered. Full-scale IQ was then estimated following the approach of Aubry and Bourdin (2018). In addition, the British Picture Vocabulary Scale (BPVS3) measure was used to assess receptive vocabulary and the Phonological Assessment Battery (PhAB, Frederickson et al., 1997; GL Assessment) was administered to assess phonological awareness at the rhyme and phoneme levels, along with rapid naming of objects and digits (RAN). The children’s amplitude rise time thresholds were estimated using 3 psychoacoustic threshold tasks based on (1) sine tones; (2) tones made from speech-shaped noise; (3) a synthetic syllable “ba”. As shown in Table 1, the children with dyslexia had significantly poorer reading and spelling skills than the control children, significantly poorer phonological skills, and significantly poorer amplitude rise time discrimination in a synthetic syllable task (discriminating differences in rise time of the syllable “ba”).

**Table 1.** Group characteristics (N=51) expressed as mean and (S.D.) for children with dyslexia and age-matched control children.

|  | Dyslexic | Age-Matched Control |
| --- | --- | --- |
| **N** | 30 | 21 |
| **Age (months)** | 110.7 (5.6) | 109.3 (5.4) |
| **WISC FSIQ** | 101.7 (10.1) | 103.6 (12.0) |
| **BAS Reading SS** | 81.0 (8.0) *** | 99.5 (6.2) |
| **BAS Reading Age in months** | 85.5 (11.0) *** | 106.5 (11.7) |
| **BAS Spelling SS** | 79.9 (7.5) *** | 97.1 (6.1) |
| **TOWRE SWE SS** | 79.5 (12.8) *** | 101.1 (7.7) |
| **TOWRE PDE SS** | 79.2 (10.9) *** | 98.0 (8.6) |
| **PhAB Rhyme SS** | 92.6 (11.7) *** | 102.4 (5.9) |
| **PhAB Phoneme SS** | 97.6 (9.8) ** | 105.1 (9.4) |
| **PhAB RAN objects SS** | 91.9 (15.0) * | 101.2 (13.0) |
| **PhAB RAN digits SS** | 85.8 (15.8) ** | 97.4 (13.3) |
| **Rise time sine tone ms** | 174.7 (81.7) | 139.7 (84.5) |
| **Rise time SSN ms** | 221.9 (56.7) | 215.0 (54.9) |
| **Rise time “ba” ms** | 101.5 (43.8)* | 70.8 (34.6) |

Note. \*\*\* $p < .001$ ; \*\* $p < .01$ ; \* $p < .05$ ; + $p = .052$ . WISC FSIQ = Wechsler Intelligence Scale for Children, estimated Full Scale IQ; BPVS = British Picture Vocabulary Scales, SS = standardised mean score = 100; BAS = British Ability Scales, SS = standardized score = 100; TOWRE SWE = Test of Word Reading Efficiency Sight Word Efficiency Scale, SS = standardized score = 100; TOWRE PDE = Phonic Decoding Efficiency Scale, SS = standardized score = 100; PhAB = Phonological Awareness Battery, SS = standardised mean score = 100; RAN = Rapid Automatized Naming; Rise time = threshold in rise time task in ms; SSN = speech-shaped noise.

### 2.2. Experimental set-up and stimuli

The children were seated in an electrically shielded soundproof room. The auditory stimuli (through earphones) were presented at a sampling rate of 44.1 kHz to the participant while EEG data were collected at a sampling rate of 1 kHz using a 128-channel EEG system (HydroCel Geodesic Sensor Net). The stimuli were natural speech presented as a 10-minute long story for children, *The Iron Man: A Children’s Story in Five Nights* by Ted Hughes. The story was read in child-directed speech and was presented in ten sections, each of which lasted about one minute followed by a 2 second gap. During the experiment, participants were instructed to listen to the speech carefully and to look at a red cross (+) shown on the screen that was in front of them.

### 2.3. Auditory Stimuli and EEG Data Pre-processing

In this study, we first obtained the broad-band envelopes by calculating the absolute value of the analytical signal of the speech stimuli. The broad-band envelopes were then filtered at frequency range of 0.5 – 8 Hz to extract the low frequency envelopes. The low frequency envelopes were used for the analyses used here.

The collected EEG data were referenced to Cz channel and then band-passed filtered into frequency range of 0.5 – 48 Hz using a zero phase FIR filter with low cutoff (−6 dB) of 0.25 Hz and high cutoff (−6 dB) of 48.25 Hz (EEGLab Toolbox; Delorme and Makeig, 2004). Bad channels were detected and interpolated through spherical interpolation (EEGLab Toolbox). The EEG data were downsampled to 100 Hz and filtered to extract delta (0.5 – 4 Hz), theta (4 – 8 Hz) and alpha (8 – 12 Hz) frequency bands. The data were further downsampled to 50 Hz to reduce the computational costs, and then epoched into the 1-minute story parts.

### 2.4. Stimulus Reconstruction Using Backward TRF Model

To investigate how accurately the children were encoding the low-frequency information in the speech signal, we used a decoding accuracy method based on the backward TRF model (Crosse et al., 2016). Given the focus of TS theory on the envelope, and given that the speech envelope carries the most critical low-frequency acoustic information aiding linguistic decoding, the speech envelope was chosen as the decoding target in our study. The TRF model was used to reconstruct the stimuli envelopes from the EEG signals at each frequency band of interest (delta, theta, alpha). The backward TRF model estimates the stimulus envelope, *s*(*t*), through the following equation:

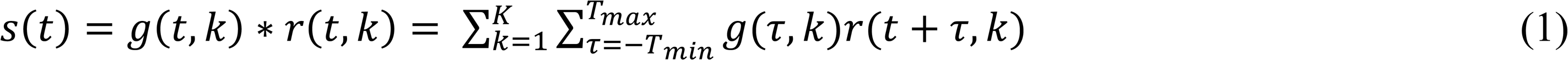

where *g*(*τ, k*) is a decoder representing the linear mapping from the neural response to the stimulus envelope for time lag *τ* and channel *k*, parameters *T*_*min*_ = 0 *ms* and *T*_*max*_ = 500 *ms* determine the minimum and maximum time lags, respectively, and *r*(*t, k*) represents the neural response at channel *k*. Accordingly, the backward model can estimate the quality of the child’s encoding of the original speech signal, as the accuracy of decoding estimated by the model provides a measure of the quality of the neural representation of the low-frequency speech information in each frequency band. To prevent overfitting and to avoid biasing the model to a specific part of the training data set, the backward model uses the Tikhonov regularization coupled with a cross-validation procedure. The validation procedure was the “leave-one-out” cross-validation (using *mTRFcrossval* function from the mTRF Toobox) in which each trial is “left out” or used for testing and the remainder are used to train the model and this procedure is repeated across all trials. Eight values were used for the ridge parameter (λ = 0.1, 1, …, 10^6^) in this procedure and the value that gave the highest average correlation score was chosen as the optimal value for the ridge parameter. This optimal value was then used to train the model.

The Pearson correlation between the estimated envelope and the actual envelope was used to estimate the performance of the model, with a higher correlation value indicating more accurate decoding by the model.

### 2.5. Between-Group Backward TRF Analysis

To model normative decoding of the low-frequency envelope information in the speech signal from the EEG signal at each frequency band, we used EEG recorded from the typically-developing (control) children. For each frequency band of interest (delta, theta, alpha), we used a randomly-selected subset of control children (*C*_*train*_ = 11) for training 11 backward TRF models. A final model was obtained by averaging across all 11 individual trained models. This final averaged model was then tested separately on the rest of the control children (*C*_*test*_ = 10) and on the 29 dyslexic children. We next computed the mean correlation values for the tested control children and the dyslexic children, resulting in a single value for each group (control, dyslexic). This procedure was then repeated 100 times, using random permutations across all the control children, in order to use different combinations of 11 versus 10 (*C*_*train*_ and *C*_*test*_) control children for training and testing the model, resulting in 100 normative (averaged) models. Accordingly, 100 correlation values were generated for the control children and 100 correlation values were generated for the dyslexic children, providing an averaged correlation value for each group. This then enabled us to assess whether there were differences in speech decoding accuracy regarding low-frequency envelope information between children with dyslexia and age-matched controls. In essence, the reconstruction accuracy for the typically-developing children was utilised as a benchmark to compare with the reconstruction accuracy obtained for the dyslexic children. Figure 1 provides a schematic diagram of the analytic approach used for the between-group backward TRF analysis.

**Figure 1.**
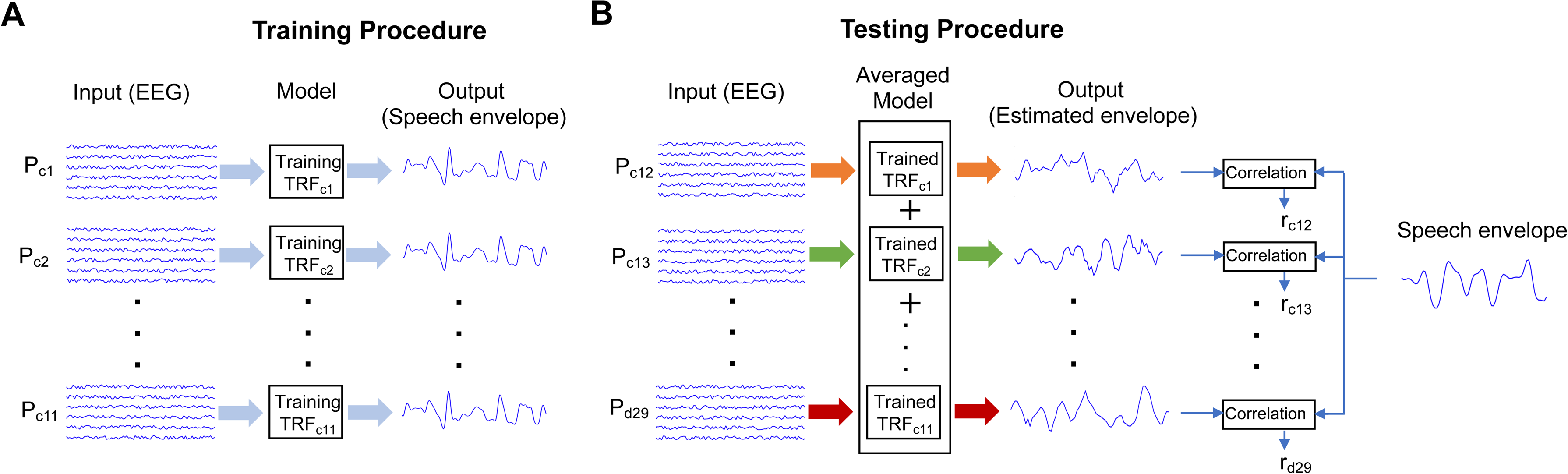
Schematic diagram for the model used for the between-group analysis. Panel A shows the procedure used to train the model and panel B shows the procedure applied to test the model. *P*_*ci*_ (*i* = 1, 2, …, 21): the *i*th control participant; *P*_*di*_ (*i* = 1, 2, …, 29): the *i*th dyslexic participant; *r*_*ci*_ (*i* = 12, 13, …, 21): decoding accuracy for the *i*th control participant; *r*_*di*_ (*i* = 1, 2, …, 29): decoding accuracy for the *i*th dyslexic participant.

### 2.6. Within-Child Backward TRF Analysis

To discover whether the perceptual experience of low-frequency envelope information in the speech signal is consistent (or stable) for any individual child, we also built a single backward TRF model for each child in the study based on the data from that child. This within-child approach enabled us to assess whether the speech-based representations developed by children with dyslexia are less consistent, “fuzzy” or “noisy” for each frequency band of interest. This was achieved by using 80% of the data (eight story sections) for each child to train the TRF model for that child, and then using the remaining data from that child (two story sections) to test the model. We calculated the Pearson correlation between the estimated speech envelope and the actual speech envelope for each child separately for each frequency band. This resulted in a single correlation score for each band for each child. These scores were then summed by group for each band and compared between groups. Figure 2 provides a schematic diagram of the analytic approach used for the within-child backward TRF analysis.

**Figure 2.**
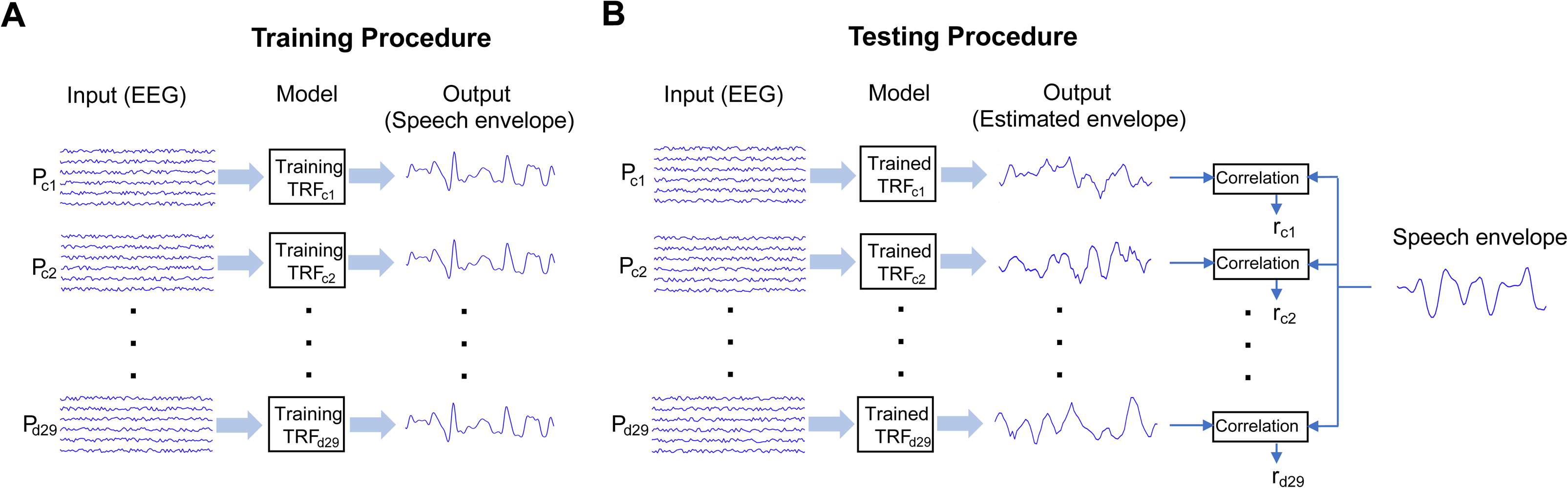
Schematic diagram for the model used for the within-child analysis. Panel A shows the procedure used to train the model and panel B illustrates the procedure applied to test the model. *P*_*ci*_ (*i* = 1, 2, …, 21): the *i*th control participant; *P*_*di*_ (*i* = 1, 2, …, 29): the *i*th dyslexic participant; *r*_*ci*_ (*i* = 1, 2, …, 21): decoding accuracy for the *i*th control participant; *r*_*di*_ (*i* = 1, 2, …, 29): decoding accuracy for the *i*th dyslexic participant.

We also applied the Wilcoxon rank sum test to compare the ridge parameter values for the within-child model between the control and dyslexic groups separately for the three frequency bands. The results showed that there was no significant difference between the two groups for all three bands (Wilcoxon rank sum test; delta, *z* = – 1.34, *p* = 0.18; theta, *z* = 0.71, *p* = 0.48; alpha, *z* = – 0.68, *p* = 0.50).

### 2.7. Computation of the Chance Level for Backward TRF

To check the statistical significance of stimulus reconstruction accuracy as estimated by the backward TRF models (both between-group and within-child), we computed null models for each frequency band of interest. To obtain the null models, we randomly selected ten control children and EEG data were permuted across different story sections for each child. We then built a single model for each child. To achieve this, 80% of the data for each child were used to train the null model for that child, and the remaining data were used to test the null model. The Pearson correlation between the estimated speech envelope and the actual speech envelope for each child separately for each frequency band was then calculated for these null models, resulting in a single correlation value for each band and for each child. We finally computed the mean reconstruction accuracy by averaging across the accuracy scores obtained for the selected children. This was done 100 times for each band, in order to calculate the probability density functions (PDF, a statistical measure that determines the probability distribution of a variable) for the null models and thereby establish chance levels. Note that the area under the PDF curve should be 1, hence for our data the smaller the range of Pearson correlations on the *x* axis, the higher the value required for the PDF on the *y* axis.

## 3. Results

### 3.1. Does the Accuracy of Speech Envelope Decoding Vary Between Control and Dyslexic Children?

To investigate whether the accuracy of speech envelope decoding estimated from EEG recorded from the dyslexic brain is different from that recorded from the typically-developing brain, the Between-Group Backward TRF analysis (see Section 2.5) was applied separately for each frequency band. To check the statistical significance of the stimulus reconstruction accuracy obtained for each frequency band by group, we first compared decoding accuracy in each band to decoding accuracy for the null models (the estimate of chance level for each band, see Section 2.7). Statistical significance by band and group is shown in Figure 3. The modelling showed that stimulus-reconstruction accuracy was significantly greater than chance in the delta band only (Figure 3A). Reconstruction accuracy in the theta and alpha bands was not statistically different from the noise values in these bands for either group (Figure 3B,C).

**Figure 3.**
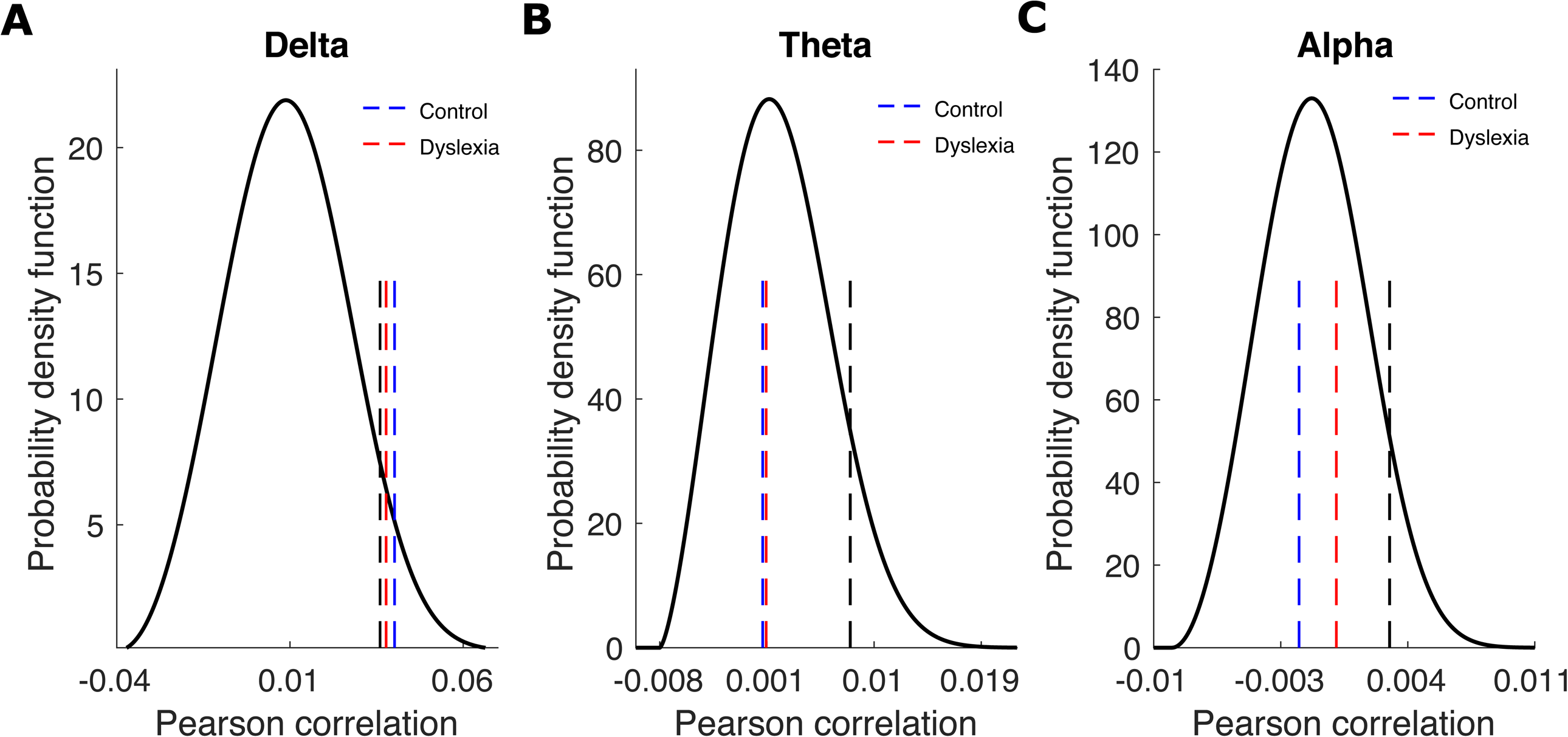
Statistical significance of reconstruction accuracy for the delta band (Panel A), theta band (Panel B) and alpha band (Panel C) for the Between-Group Analyses (Blue, control children; Red, dyslexic children). The probability density functions of the null models are shown by the black curves, please note that the scales for the three bands are different. The dashed black lines denote the correlation values corresponding to statistical significance (*p* = 0.05). The blue and red dashed lines show the mean correlation values obtained for the control and dyslexic groups, respectively.

To compare the accuracy of decoding between the control and dyslexic groups in the delta band, we applied the Wilcoxon rank sum test. This showed that reconstruction accuracy for the control group in the delta band as derived from 100 TRF models was significantly greater than reconstruction accuracy for the dyslexic group (Wilcoxon rank sum test, *z* = 3.72, *p* = 0.0002). This difference in speech envelope decoding suggests that control brains encode a more accurate neural representation of the speech envelope in the delta band compared to dyslexic brains, as shown in Figure 4.

**Figure 4.**
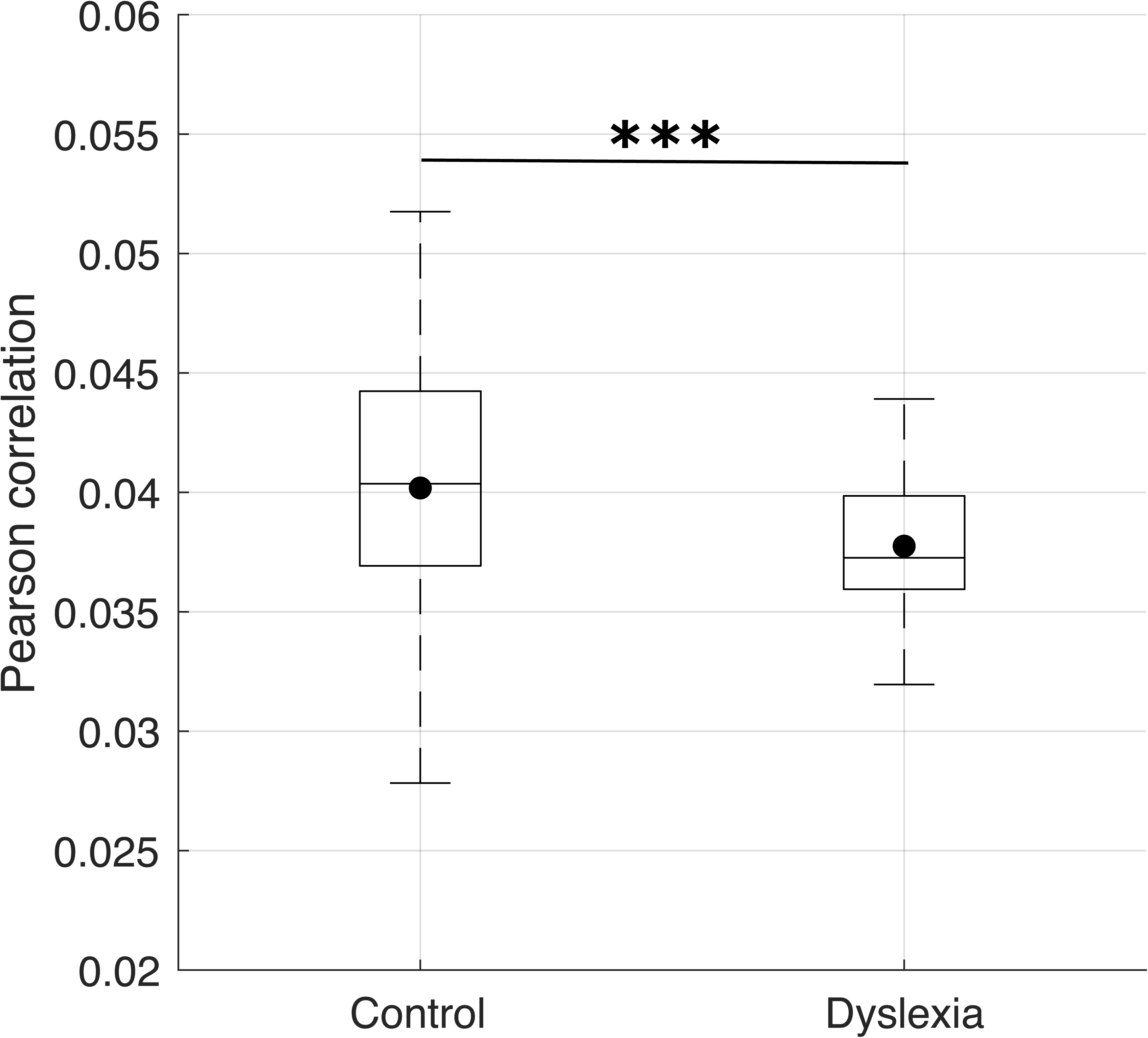
Stimulus-reconstruction accuracy for control versus dyslexic children in the delta band summarising correlation values from 100 backward TRF models. The black circles show the mean accuracy scores. On each box, the central line indicates the median, and the bottom and top edges of the box denote the 25th and 75th percentiles, respectively. The whiskers also extend to the most extreme data points. *** denotes *p* < 0.001.

### 3.2. Do Children with Dyslexia Develop “Noisy” Representations of Low-Frequency Information in Speech?

To explore whether children with dyslexia have “noisy” or “fuzzy” representations of the low-frequency envelope information in the speech signal, the Within-Child Backward TRF analysis method (see Section 2.6) was employed separately for each frequency band. The statistical significance of the stimulus reconstruction accuracy obtained for each frequency band for the within-child analyses was computed by considering chance level for that band (the null models, see Section 2.7). Comparisons with the null models showed that stimulus-reconstruction accuracy was above chance-levels (alpha = 0.05) for both groups in all three frequency bands (Figure 5).

**Figure 5.**
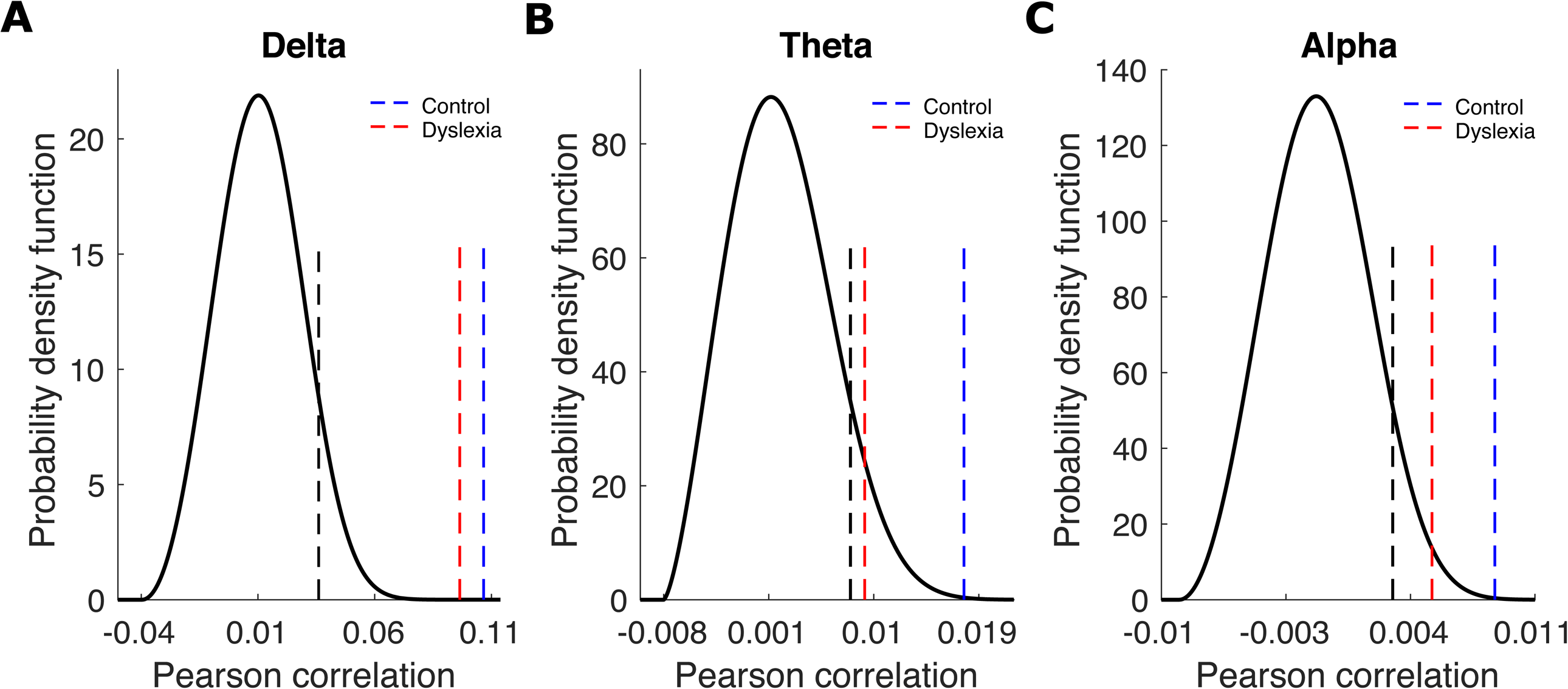
Statistical significance of within-child stimulus reconstruction accuracy for the delta band (Panel A), theta band (Panel B) and alpha band (Panel C) summarised by group (Blue, control children; Red, dyslexic children). The probability density functions of the null models are again shown by the black curves. The dashed black lines denote the correlation values corresponding to statistical significance (*p* = 0.05). The blue and red dashed lines show the mean correlation values obtained from the within-child analysis for control and dyslexic groups, respectively.

Figure 6 shows the box plots for stimulus reconstruction accuracies for the control and dyslexic groups based on the backward TRF models from individual children (shown as dots) by band, separated by group. To compare reconstruction accuracy between the two groups, we employed the Wilcoxon rank sum test. The results by band showed that there was no difference between the two groups in any band (delta band, Figure 6A; Wilcoxon rank sum test, *z* = – 0.59, *p* = 0.55; theta band, Figure 6B; Wilcoxon rank sum test, *z* = −1.30, *p* = 0.19; alpha band, Figure 6C; Wilcoxon rank sum test, *z* = – 0.81, *p* = 0.42). We also applied Bayesian factor analysis to quantify the strength of the evidence for the alternative model. Bayesian factor analysis estimates the strength of the evidence for the alternate hypothesis H_1_ that the groups differ in their stimulus reconstruction accuracy over the null hypothesis H_0_ that they do not. The results indicated greater evidence in favour of the null model for all three bands (Bayesian factor analysis; delta band, *BF10* = 0.2; theta band, *BF10* = 0.9; alpha band, *BF10* = 0.3). Accordingly, as shown in Figure 6, the backward TRF model can reconstruct the stimulus envelopes for each individual child in each band consistently from that child’s EEG data. Furthermore, this envelope information is reconstructed above chance levels for each individual child in all three bands, in contrast to the reconstruction consistency achieved for the between-group analysis, in which decoding accuracy was only above chance for both groups of children in the delta band condition (Figure 3). The within-child group comparisons (Figure 6) show that the consistency of decoding accuracy is always above chance, and is not statistically different irrespective of whether the child is a child with dyslexia or a control child. The modelling indicates that the neural representation of the low-frequency envelope information in the speech signal for each child is consistent for that child, providing them with a stable percept of this aspect of speech.

**Figure 6.**
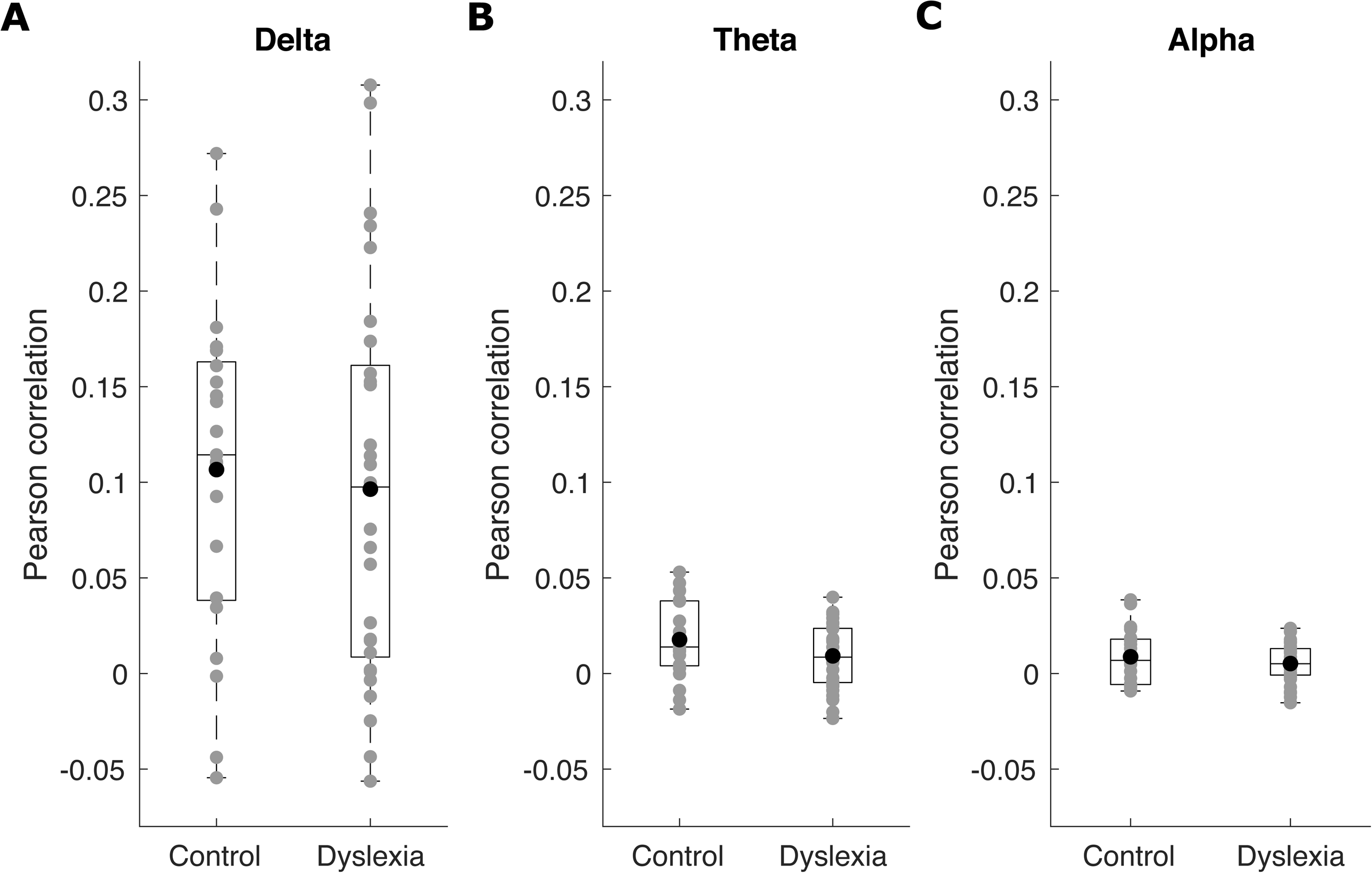
Stimulus-reconstruction accuracy computed for each individual control and dyslexic child in the delta (Panel A), theta (Panel B), and alpha (Panel C) bands in the within-child analysis. The grey disks denote individual children’s reconstruction accuracy scores, and the black circles show the mean accuracy scores for each band and each group.

## 4. Discussion

Here we employed backward TRF models to estimate stimulus reconstruction accuracy of bands of low-frequency acoustic information in the speech signal (delta- and theta-band speech envelope information) along with alpha-band speech envelope information (control band) in children with and without dyslexia matched for age. Two different group comparison approaches (between-group and within-child) were employed to investigate whether the speech-based representations for low-frequency envelope information being developed by the dyslexic brain were “noisy” or “normal”. If decoding accuracy varies by group for the between-group TRF models, then the speech-based representations of low-frequency acoustic information developed by children with developmental dyslexia cannot be considered “normal”. However, if decoding accuracy is equal irrespective of group for within-child TRF models, then the speech-based representations of low-frequency envelope information developed by children with developmental dyslexia cannot be considered “noisy”. The modelling suggested that the dyslexic brain does not develop “noisy” representations of low-frequency envelope information, but neither are these representations “normal”. The between-group models showed that the speech-based representations developed by children with dyslexia were *less accurate* than the speech-based representations developed by children without dyslexia regarding low frequency envelope information. These differences were restricted in the current study to the delta band (Figure 4). The between-group modelling generated lower decoding accuracies than the within-child modelling, as can be seen by comparing the correlation values in Figures 4 and 6. Between-group decoding accuracy for the theta and alpha bands did not exceed chance levels as estimated by the null models. Therefore, the conservative conclusion is that encoding of low-frequency speech information in the delta band is atypical in 9-year-old children with developmental dyslexia.

At the level of an individual child, however, the modelling did not support the classical theoretical view (including our own earlier view, see Swan & Goswami, 1997) that the phonological representations of speech developed by children with dyslexia are “fuzzy”, “noisy” or “imprecise” (see also Elbro et al., 1998; Snowling, 2000). Although we only decoded the low-frequency envelope information in speech, the quality of the speech-based representations for each child as indexed by the neural decoding method were of similar consistency whether the child was dyslexic or not, for all frequency bands explored. This finding suggests that even if acoustic information is weighted differently in the speech-based representations developed by children with dyslexia, the experience of speech processing regarding low-frequency envelope information is perceptually stable for the children themselves. Nevertheless, a linear decoding approach to stimulus reconstruction is only one method for assessing whether neural stimulus representations of low-frequency envelope information are “noisy” or not, and converging neural methods are required. Furthermore, the question of whether other features of the speech signal such as voicing may be represented in a “noisy” manner by the dyslexic brain is not addressed by our data.

The delta-band between-group difference in decoding accuracy found here was predicted by TS theory (Goswami, 2011, 2019), which provided the conceptual framework for the current study. TS theory has proposed that dyslexic difficulties in discriminating amplitude envelope rise times, difficulties found for dyslexic children in 7 languages to date (see Goswami, 2015, for a review), are associated with atypical neural encoding of acoustic information within the amplitude envelope < 10Hz, related to perceptual impairments in processing speech rhythm. Rise times are one neural trigger for oscillatory phase-resetting, an automatic process that aids multi-timescale speech-brain cortical tracking in adults (Giraud & Poeppel, 2012; Doelling et al., 2014; Lizarazu et al., 2021). A recent MEG study with adults with dyslexia has shown that rise times do not provide an efficient phase-resetting mechanism in the dyslexic brain during natural speech listening (Lizarazu et al., 2021). The between-group data presented here suggest that this inefficient phase-resetting may particularly affect the encoding of speech envelope information in the delta band during childhood. This delta-band information is a critical feature of the speech signal.

The current study has several limitations. As already noted, the quality of dyslexic children’s speech-based representations for features other than the envelope is not addressed by our backward decoding method. Further, although the backward TRF model has been used as the most popular model for decoding stimulus information from neural responses, it has some general limitations. Firstly, the model assumes a linear relationship between the input stimuli and the neural responses. Secondly, the performance of model when generalising to unseen testing data sets depends on the estimate of large number of unknown parameters and on the size of the data used for training the model. It should be noted that the forward TRF model also has these two latter limitations. However, many adult studies focus on backward (decoding) models, as they have several advantages over forward (encoding) models, as described by Crosse et al. (2016).

In conclusion, the current study converges with the three preceding studies of children with dyslexia that measured speech-based representations directly using neural methods (Power et al., 2016; Di Liberto et al., 2018; Destoky et al., 2020). These studies all reported that children with dyslexia show atypical encoding of low-frequency information in the speech amplitude envelope. Together with the current study, these studies suggest that the speech-based representations developed by children with dyslexia for low-frequency envelope information <10 Hz are not “normal”, supporting TS theory. Children with dyslexia encode less accurate prosodic (delta band) speech-based information in their phonological representations for words in comparison to typically-developing children. TS theory proposes that these differences in encoding low-frequency envelope information affect phonological awareness at all linguistic levels via the linguistic hierarchy, which is governed by the prosodic level, and that this then impacts the process of learning to read and spell. At the same time, the current study contributes a novel child-by-child neural decoding approach that reveals that the perceptual world of the dyslexic child regarding low-frequency envelope information in speech is a consistent one. At the level of the individual child, the speech-based representations of low-frequency envelope information developed by children with dyslexia are not “fuzzy” or “noisy”. This finding contributes to a long-standing debate in the psycholinguistic literature regarding whether the phonological ‘deficit’ that characterises individuals with dyslexia across languages stems from differences in phonological representation or in phonological access (e.g., Adlard and Hazan, 1998; Elbro et al., 1998; Manis et al., 1997; Mody, Studdert-Kennedy, and Brady, 1997; Serniclaes et al., 2004; Bogliotti et al., 2008; Snowling, 2000; Ramus & Svenkovits, 2008). The neural decoding method employed here suggests that differences in phonological representation are indeed present for low-frequency speech information. This conclusion accords well with the linguistic data gathered by recent longitudinal studies of infants at family risk for developmental dyslexia, who show amplitude rise time discrimination difficulties from age 10 months as well as poorer early word learning and poorer phonological development (Kalashnikov et al., 2018, 2019a,b, 2020). Nevertheless, the strongest test of this conclusion about phonological representations would be a demonstration that speech production is also atypical in dyslexia regarding the specification of speech prosody. If speech prosody is not encoded accurately in the dyslexic brain, yet this atypical encoding is consistent and perceptually stable for the child, then speech output of prosodic information should also be affected.

## Acknowledgment

The research was funded by a grant awarded to UG by the Foundation Botnar (project number: 6064) and a donation from the Yidan Prize Foundation. The sponsors played no role in the study design, data interpretation or writing of the report. The authors would like to thank all the families and schools involved in the study.

## References

Adlard, A., & Hazan, V. (1998). Speech perception in children with specific reading difficulties (dyslexia). The Quarterly Journal of Experimental Psychology Section A, 51(1), 153–177. https://doi.org/10.1080/713755750.

Aubry, A., & Bourdin, B. (2018). Short Forms of Wechsler scales assessing the intellectually gifted children using simulation data. Frontiers in psychology, 9, 830. https://doi.org/10.3389/fpsyg.2018.00830.

Bogliotti, C., Serniclaes, W., Messaoud-Galusi, S., & Sprenger-Charolles, L. (2008). Discrimination of speech sounds by children with dyslexia: Comparisons with chronological age and reading level controls. Journal of experimental child psychology, 101(2), 137–155. https://doi.org/10.1016/j.jecp.2008.03.006.

Catts, H. W. (1993). The relationship between speech-language impairments and reading disabilities. Journal of Speech, Language, and Hearing Research, 36(5), 948–958. https://doi.org/10.1044/jshr.3605.948.

Crosse, M. J., Di Liberto, G. M., Bednar, A., & Lalor, E. C. (2016). The multivariate temporal response function (mTRF) toolbox: a MATLAB toolbox for relating neural signals to continuous stimuli. Frontiers in human neuroscience, 10, 604. https://doi.org/10.3389/fnhum.2016.00604.

Delorme, A., & Makeig, S. (2004). EEGLAB: an open-source toolbox for analysis of singletrial EEG dynamics including independent component analysis. Journal of neuroscience methods, 134, 9–21. https://doi.org/10.1016/j.jneumeth.2003.10.009.

Destoky, F., Bertels, J., Niesen, M., Wens, V., Vander Ghinst, M., Leybaert, J., Lallier, M., Ince, R. A., Gross, J., De Tiège, X., & Bourguignon, M. (2020). Cortical tracking of speech in noise accounts for reading strategies in children. PLoS biology, 18(8), p.e3000840. https://doi.org/10.1371/journal.pbio.3000840.

Di Liberto, G. M., O’Sullivan, J. A., & Lalor, E. C. (2015). Low-frequency cortical entrainment to speech reflects phoneme-level processing. Current Biology, 25(19), 2457–2465. https://doi.org/10.1016/j.cub.2015.08.030.

Di Liberto, G. M., Peter, V., Kalashnikova, M., Goswami, U., Burnham, D., & Lalor, E. C. (2018). Atypical cortical entrainment to speech in the right hemisphere underpins phonemic deficits in dyslexia. NeuroImage, 175, 70–79. https://doi.org/10.1016/j.neuroimage.2018.03.072.

Dimitrijevic, A., Smith, M. L., Kadis, D. S. & Moore, D. R. (2017). Cortical alpha oscillations predict speech intelligibility. Frontiers in human neuroscience, 11, 88. https://doi.org/10.3389/fnhum.2017.00088.

Ding, N. and Simon, J. Z. (2014). Cortical entrainment to continuous speech: functional roles and interpretations. Frontiers in human neuroscience, 8, 311. https://doi.org/10.3389/fnhum.2014.00311.

Doelling, K. B., Arnal, L. H., Ghitza, O., & Poeppel, D. (2014). Acoustic landmarks drive delta-theta oscillations to enable speech comprehension by facilitating perceptual parsing. NeuroImage, 85, 761–768. https://doi.org/10.1016/j.neuroimage.2013.06.035.

Elbro, C., Borstrøm, I., & Petersen, D. K. (1998). Predicting dyslexia from kindergarten: The importance of distinctness of phonological representations of lexical items. Reading research quarterly, 33(1), 36–60. https://doi.org/10.1598/RRQ.33.1.3.

Elliott, C. D., Smith, P., & McCullogh, K. (1996). British Ability Scales (2nd ed.). Windsor, UK: NFER-Nelson.

Ehri, L. C., & Wilce, L. S. (1980). The influence of orthography on readers’ conceptualization of the phonemic structure of words. Applied Psycholinguistics, 1(4), 371–385. https://doi.org/10.1017/S0142716400009802.

Frederickson, N., Frith, U., & Reason, R. (1997). Phonological Assessment Battery: Standardised Edition. NFER-Nelson.

Giraud, A.L., & Poeppel, D. (2012). Cortical oscillations and speech processing: Emerging computational principles and operations. Nature Neuroscience, 15(4), 511–517. https://doi.org/10.1038/nn.3063.

Goswami, U. (2011). A temporal sampling framework for developmental dyslexia. Trends in Cognitive Sciences, 15(1), 3–10. https://doi.org/10.1016/j.tics.2010.10.001.

Goswami, U. (2015). Sensory theories of developmental dyslexia: Three challenges for research. Nature Reviews Neuroscience, 16(1), 43–54. https://doi.org/10.1038/nrn3836.

Goswami, U. (2019). Speech rhythm and language acquisition: An amplitude modulation phase hierarchy perspective. Annals of the New York Academy of Sciences, e14137. https://doi.org/10.1111/nyas.14137.

Goswami, U., 2022. Theories of Developmental Dyslexia. To appear in M. Skeide (Ed.), The Cambridge Handbook of Dyslexia and Dyscalculia. Cambridge, UK: Cambridge University Press.

Goswami, U., Fosker, T., Huss, M., Mead, N., & Szücs, D. (2011). Rise Time and Formant Transition Duration in the Discrimination of Speech Sounds: The Ba-Wa Distinction in Developmental Dyslexia. Developmental Science, 14, 34–43. https://doi.org/10.1111/j.1467-7687.2010.00955.x.

Guttorm, T. K., Leppänen, P. H., Poikkeus, A. M., Eklund, K. M., Lyytinen, P., & Lyytinen, H. (2005). Brain event-related potentials (ERPs) measured at birth predict later language development in children with and without familial risk for dyslexia. Cortex, 41(3), 291–303. https://doi.org/10.1016/S0010-9452(08)70267-3.

Guttorm, T. K., Leppänen, P. H., Hämäläinen, J. A., Eklund, K. M., & Lyytinen, H. J. (2010). Newborn event-related potentials predict poorer pre-reading skills in children at risk for dyslexia. Journal of learning disabilities, 43(5), 391–401. https://doi.org/10.1177/0022219409345005.

Kalashnikova, M., Goswami, U., & Burnham, D. (2018). Mothers speak differently to infants at-risk for dyslexia. Developmental Science, 21(1), 1–15. https://doi.org/10.1111/desc.12487.

Kalashnikova, M., Goswami, U., & Burnham, D. (2019a). Delayed development of phonological constancy in toddlers at family risk for dyslexia. Infant Behavior and Development, 57, 101327. https://doi.org/10.1016/j.infbeh.2019.101327.

Kalashnikova, M., Goswami, U., & Burnham, D. (2019b). Sensitivity to amplitude envelope rise time in infancy and vocabulary development at 3 years: A significant relationship. Developmental Science, 22(6), 1–9. https://doi.org/10.1111/desc.12836.

Kalashnikova, M., Goswami, U., & Burnham, D. (2020). Novel word learning deficits in infants at family risk for dyslexia. Dyslexia, e1649. https://doi.org/10.1002/dys.1649.

Keshavarzi, M., Kegler, M., Kadir, S. & Reichenbach, T. (2020). Transcranial alternating current stimulation in the theta band but not in the delta band modulates the comprehension of naturalistic speech in noise. NeuroImage, 210, 116557. https://doi.org/10.1016/j.neuroimage.2020.116557.

Keshavarzi, M. & Reichenbach, T. (2020). Transcranial alternating current stimulation with the theta-band portion of the temporally-aligned speech envelope improves speech-in-noise comprehension. Frontiers in Human Neuroscience, 14, 187. https://doi.org/10.3389/fnhum.2020.00187.

Keshavarzi, M., Mandke, K., Macfarlane, A., Parvez, L., Gabrielczyk F., Wilson, A., & Goswami, U. (2022). Atypical Delta-band Phase Consistency and Atypical Preferred Phase in Children with Dyslexia during Neural Entrainment to Rhythmic Audio-Visual Speech. NeuroImage: Clinical, 103054. https://doi.org/10.1016/j.nicl.2022.103054.

Leppänen, P. H., Hämäläinen, J. A., Salminen, H. K., Eklund, K. M., Guttorm, T. K., Lohvansuu, K., Puolakanaho, A., & Lyytinen, H. (2010). Newborn brain event-related potentials revealing atypical processing of sound frequency and the subsequent association with later literacy skills in children with familial dyslexia. Cortex, 46(10), 1362–1376. https://doi.org/10.1016/j.cortex.2010.06.003.

Lizarazu, M., Lallier, M., Bourguignon, M., Carreiras, M., & Molinaro, N. (2021). Impaired neural response to speech edges in dyslexia. Cortex, 135, 207–218. https://doi.org/10.1016/j.cortex.2020.09.033.

Manis, F. R., McBride-Chang, C., Seidenberg, M. S., Keating, P., Doi, L. M., Munson, B., & Petersen, A. (1997). Are speech perception deficits associated with developmental dyslexia?. Journal of experimental child psychology, 66(2), 211–235. https://doi.org/10.1006/jecp.1997.2383.

Mody, M., Studdert-Kennedy, M., & Brady, S. (1997). Speech perception deficits in poor readers: Auditory processing or phonological coding?. Journal of experimental child psychology, 64(2), 199–231. https://doi.org/10.1006/jecp.1996.2343.

Power, A. J., Colling, L. J., Mead, N., Barnes, L., & Goswami, U. (2016). Neural encoding of the speech envelope by children with developmental dyslexia. Brain and Language, 160, 1–10. https://doi.org/10.1016/j.bandl.2016.06.006.

Power, A. J., Mead, N., Barnes, L., & Goswami, U. (2012). Neural entrainment to rhythmically presented auditory, visual, and audio-visual speech in children. Frontiers in Psychology, 3, 216. https://doi.org/10.3389/fpsyg.2012.00216.

Ramus, F., & Szenkovits, G. (2008). What phonological deficit?. Quarterly journal of experimental psychology, 61(1), 129–141. https://doi.org/10.1080/17470210701508822.

Saffran, J.R. (2001). Words in a sea of sounds: The output of infant statistical learning. Cognition, 81(2), 149–169. https://doi.org/10.1016/S0010-0277(01)00132-9.

Serniclaes, W., Van Heghe, S., Mousty, P., Carré, R., & Sprenger-Charolles, L. (2004). Allophonic mode of speech perception in dyslexia. Journal of experimental child psychology, 87(4), 336–361. https://doi.org/10.1016/j.jecp.2004.02.001.

Serniclaes, W., & Seck M. (2018). Enhanced Sensitivity to Subphonemic Segments in Dyslexia: A New Instance of Allophonic Perception. Brain Sciences, 8(4), 54. https://doi.org/10.3390/brainsci8040054

Snowling, M.J. (2000). Dyslexia. Blackwell Publishers, Oxford.

Stackhouse, J., & Wells, B. (1997). Children’s Speech and Literacy Difficulties, Book1: A Psycholinguistic Framework, 9. Wiley-Blackwell.

Stanovich, K. E., & Siegel, L. S. (1994). Phenotypic performance profile of children with reading disabilities: A regression-based test of the phonological-core variable-difference model. Journal of educational psychology, 86(1), 24. https://doi.org/10.1037/0022-0663.86.1.24.

Stein, J., & Walsh, V. (1997). To see but not to read; the magnocellular theory of dyslexia. Trends in neurosciences, 20(4), 147–152. https://doi.org/10.1016/S0166-2236(96)01005-3.

Strauß, A., Wöstmann, M. & Obleser, J. (2014). Cortical alpha oscillations as a tool for auditory selective inhibition. Frontiers in human neuroscience, 8, 350. https://doi.org/10.3389/fnhum.2014.00350.

Swan, D., & Goswami, U. (1997). Phonological awareness deficits in developmental dyslexia and the phonological representations hypothesis. Journal of experimental child psychology, 66(1), 18–41. https://doi.org/10.1006/jecp.1997.2375.

Torgesen, J., Wagner, R. K., & Rashotte, C. (1999). Test of Word Reading Efficiency (TOWRE). Austin, TX: Pro-Ed.

van Zuijen, T. L., Plakas, A., Maassen, B. A., Maurits, N. M., & van der Leij, A. (2013). Infant ERPs separate children at risk of dyslexia who become good readers from those who become poor readers. Developmental science, 16(4), 554–563. https://doi.org/10.1111/desc.12049.

Wechsler, D. (2016). Wechsler Intelligence Scale for Children, 5th Edn. London, UK: Pearson Assessment.

Ziegler, J. C., & Ferrand, L. (1998). Orthography shapes the perception of speech: The consistency effect in auditory word recognition. Psychonomic Bulletin & Review, 5(4), 683–689. https://doi.org/10.3758/BF03208845.

Ziegler, J. C., & Goswami, U. (2005). Reading acquisition, developmental dyslexia, and skilled reading across languages: a psycholinguistic grain size theory. Psychological bulletin, 131, 3. https://doi.org/10.1037/0033-2909.131.1.3.

